# In-depth proteomic characterization of *Schistosoma haematobium*: towards the development of new tools for elimination

**DOI:** 10.1101/486662

**Authors:** Javier Sotillo, Mark S. Pearson, Luke Becker, Gebeyaw G. Mekonnen, Abena S. Amoah, Govert van Dam, Paul L.A.M. Corstjens, Takafira Mduluza, Francisca Mutapi, Alex Loukas

## Abstract

**Background:** Schistosomiasis is a neglected disease affecting hundreds of millions worldwide. Of the three main species affecting humans, *Schistosoma haematobium* is the most common, and is the leading cause of urogenital schistosomiasis. *S*. *haematobium* infection can cause different urogential clinical complications, particularly in the bladder, and furthermore, this parasite has been strongly linked with squamous cell carcinoma. A comprehensive analysis of the molecular composition of its different proteomes will contribute to developing new tools against this devastating disease.

**Methods and Findings:** By combining a comprehensive protein fractionation approach consisting of OFFGEL electrophoresis with high-throughput mass spectrometry, we have performed the first in-depth characterisation of the different discrete proteomes of *S*. *haematobium* that are predicted to interact with human host tissues, including the secreted and tegumental proteomes of adult flukes and secreted and soluble egg proteomes. A total of 662, 239, 210 and 138 proteins were found in the adult tegument, adult secreted, soluble egg and secreted egg proteomes, respectively. In addition, we probed these distinct proteomes with urine to assess urinary antibody responses from naturally infected human subjects with different infection intensities, and identified adult fluke secreted and tegument extracts as being the best predictors of infection.

**Conclusion:** We provide a comprehensive dataset of proteins from the adult and egg stages of *S*. *haematobium* and highlight their utility as diagnostic markers of infection intensity for the development of novel tools to control this important neglected tropical disease.

**Author Summary:** Schistosomiasis is a neglected tropical disease affecting millions of people worldwide. Of the main three species affecting humans, *Schistosoma haematobium* is the most common, and is the leading cause of urogenital schistosomiasis. This parasite can cause a range of clinical complications associated with bladder pathogenesis, including squamous cell carcinoma as well as genital malignancy in women. Herein, we have performed the first comprehensive characterisation of the proteins implicated in host-parasite interactions (secreted and surface proteins from the adult flukes and secreted and soluble egg proteins) in order to advance our understanding of the parasite’s biology. Furthermore, we have characterised the different antibody responses in urine from infected human subjects from an endemic area presenting different infection intensities. The data obtained in this study can be used as a first step towards the development of novel tools for the control of urogenital schistosomiasis.

## Introduction

Schistosomiasis is a neglected tropical and debilitating disease caused by different trematodes from the genus *Schistosoma* [1]. It affects over 250 million people worldwide, particularly in developing and tropical regions [2–4]. Despite widespread use of the anthelmintic praziquantel in mass drug administration programs over the last 30 years [5], this parasitic infection still causes a loss of 1.9 million disability-adjusted life years (DALYs) [6], and this number could be greater if morbidity associated with asymptomatic infections was included in the calculations [7]. Of the 6 species affecting humans, *S*. *haematobium* is the most common, causing urogenital schistosomiasis in over 100 million people [1], although it is considered the neglected schistosome since the amount of “omics*”* information is scarce compared to other schistosomes and the difficulty in maintaining the parasite in an animal model [8, 9]. *S*. *haematobium* infection has been reported in 54 countries [10], particularly in sub-Saharan Africa and the Middle East [2, 3]. Furthermore, an outbreak of urogenital schistosomiasis was observed in Corsica (France) [11], although this parasite has recently been shown to be a hybrid between *S*. *haematobium* and *Schistosoma bovis*. [12].

The clinical complications associated with urogenital schistosomiasis are linked among others, to bladder pathology [13]. The association between squamous cell carcinoma and *S*. *haematobium* infection is undisputed [14]; indeed, this blood fluke has been classified as a Class I carcinogen by the International Agency for Research on Cancer (IARC) [15]. Once established in the mesenteric veins surrounding the bladder, *S*. *haematobium* adult females start laying eggs that pass through the bladder epithelium to be secreted in the urine. However, some eggs get trapped in the bladder wall causing a chronic local inflammation that will develop into a granuloma accompanied by relentless cell proliferation and ultimately in some patients, bladder cancer [16] as well as genital malignancy in women [17, 18]. The initial inflammatory response is thought to be a reaction to mechanical damage caused by passing of eggs through the urothelium (a multilayered epithelium that lines most of the urogenital tract) [13]; however, proteins secreted by parasite eggs have also been shown to increase cell proliferation and angiogenesis [13]. One of the most abundant proteins secreted by *S*. *haematobium* eggs is the IPSE, or interleukin-4 inducing principle from *Schistosoma mansoni* eggs. IPSE induces cell proliferation and angiogenesis [13, 19], and an IPSE homologue from *S*. *haematobium* has been shown to attenuate inflammation by stimulating release of IL-4 [19, 20].

Despite the clear role of soluble egg proteins in the development of granulomas and a carcinogenic environment, the protein composition of this or other *S*. *haematobium* tissues/organs has not been fully characterised, although a few studies have identified several immunogenic proteins [21]. In contrast, different proteomes from the related species *S*. *mansoni, S*. *japonicum* and even from the cattle-infecting *S*. *bovis* have been well characterised [22–26], which has allowed for the identification of novel vaccine and diagnostic antigen candidates. Indeed, different tegumental and secreted proteins are potential vaccine candidates and some have entered or completed Phase I and II clinical trials [27]. The most progressed vaccine candidate for *S*. *haematobium* infection is glutathione S-transferase (Sh28GST), which completed Phase III clinical testing in 2012 [28] but the results have yet to be published [29].

Characterising the molecular interface of host-parasite interactions in *S*. *haematobium* infection is crucial for (i) a better understanding of the parasite’s biology and, (ii) for the development of new tools for control and diagnosis. In the present study, we provide the first in-depth identification of secreted and surface proteomes of *S*. *haematobium*. We also characterise the antibody responses in urine from infected human subjects from an endemic area presenting with different infection intensities as a first step towards the development of novel diagnostic tools against this devastating disease.

## Experimental procedures

### Ethics

The collection of urine from individuals from Zimbabwe was approved by the Medical Research Council of Zimbabwe; Approval MRCZ/A/1710.

All experimental procedures involving animals reported in the study were approved by the James Cook University (JCU) animal ethics committee (ethics approval number A2391). The study protocols were in accordance with the 2007 Australian Code of Practice for the Care and Use of Animals for Scientific Purposes and the 2001 Queensland Animal Care and Protection Act.

### Human Schistosoma haematobium parasitology

Urine samples from naturally infected individuals were collected on three consecutive days for parasitological examinations. *S*. *haematobium* infection was detected by microscopic examination of the parasite eggs in 10ml urine, processed using the standard urine filtration method [30]. For each participant, infection intensity was expressed as the arithmetic mean of egg counts per 10 mL urine of samples collected on consecutive days. Urines were stratified according to WHO classification as having either a high (>50 eggs/10 ml of urine), medium (11-49 eggs/10ml of urine) or low (0.3-10 eggs/10 ml of urine) level of infection [31]. Egg negative urines (0 eggs/10 ml of urine) were tested for the presence of circulating anodic antigen (CAA – a more sensitive diagnostic test than microscopic detection of eggs in urine) using the UCAA2000 (wet format) as described previously [32] to confirm the presence or absence of infection.

### Parasite material

*S*. *haematobium*-infected *Bulinus truncatus* snails were provided by the National Institute of Allergy and Infectious Diseases (NIAID) Schistosomiasis Resource Center for distribution through BEI Resources, NIAID, National Institutes of Health (NIH), USA: *S*. *haematobium*, Egyptian strain, exposed *B*. *truncatus* subsp. *truncatus*, NR-21965. Snails were removed from their tank, washed with water and transferred to a petri dish without light or water at 27°C for 90 minutes. They were then rinsed again and water was added to an approx. depth of 3-5 mm. Snails were then placed under direct light maintaining a temperature of 28-30°C for 1-2 hours. The water was then transferred to a new petri dish through a sieve with 20 µm pore size to concentrate cercariae. Water was transferred from the snails to the sieve every 20 min three more times while cercariae continued to be shed. Mice (6 week-old Balb/c) were infected with 1,000 cercariae by tail penetration and adult worms were recovered by vascular perfusion at 16 weeks p.i. [33]. Eggs from livers of perfused mice were isolated according to the method of Dalton et al. [34] to obtain highly purified ova that were free of host debris.

### Isolation of adult excretory/secretory products

Two hundred freshly perfused adult fluke pairs were washed 3× in serum-free Basch media supplemented with 4× antibiotic/antimycotic (10,000 units/mL of penicillin, 10,000 µg/mL of streptomycin, and 25 µg/mL of Amphotericin B) (AA) (Thermo Fisher Scientific, USA) [35] followed by incubation in the same medium at 37°C, 5% CO_2_ at a density of ∼50 fluke pairs in 4 mL of medium for 7 days. Culture medium was changed after 4 hours and discarded to minimize the presence of host contaminants from regurgitating flukes. Media was subsequently changed every day and dead flukes removed from the plate to avoid contamination with somatic proteins. The excretory/secretory (ES) material was collected each day, centrifuged at 500 *g*, 2,000 *g* and 4,000 *g* to remove parasite debris, buffer exchanged in PBS, concentrated using a 10 kDa spin concentrator (Merck Millipore, USA), protein quantified by BCA (Thermo Fisher Scientific, USA), aliquoted and stored at −80°C until use.

### Tegument extraction

Extraction of tegument proteins from adult flukes was performed using the freeze/thaw/vortex technique as described previously [22, 36]. Briefly, two batches of 50 freshly perfused adult fluke pairs were washed 3× in PBS, frozen at −80°C, thawed on ice, washed in TBS (10 mM Tris/HCl, 0.84% NaCl, pH 7.4) and incubated for 5 min on ice in 10 mM Tris/HCl, pH 7.4. Each sample was vortexed for 5 × 1 s bursts, the tegument extract pelleted at 1,000 *g* for 30 min and solubilized three times in 200 μl of solubilizing solution (0.1% (w/v) SDS, 1.0% (v/v) Triton X-100 in 40 mM Tris, pH 7.4) with pelleting at 15,000 *g* between each wash. The washes were combined and buffer exchanged and concentrated as described above.

### Egg excretory/secretory products and soluble egg antigen

Purified eggs were cultured in serum-free Basch medium supplemented with 4× AA (50,000 eggs/5 ml) for 72 hours at 37°C with 5% CO_2_. Medium containing egg ES products was harvested every 24 hours and processed as described for adult ES. Approximately 400,000 eggs were used for ES generation. To obtain SEA, freshly isolated eggs in PBS (100,000/ml) were homogenized in a hand-held Potter-Elvehjem glass homogeniser (15 ml capacity), centrifuged at 200 *g* for 20 min at 4°C and then the supernatant centrifuged at 100,000 *g* for 90 min at 4°C [33]. The supernatant was then processed as described for adult ES.

### OFFGEL electrophoresis

OFFGEL fractionation was performed for adult ES, adult tegument, adult somatic and egg ES samples as described by Sotillo et al. [37]. A total of 100 µg of protein was resuspended in 50 mM NH_4_CO_3_, 20 mM DTT and incubated at 65°C for 1 h. Alkylation was then achieved by adding iodoacetamide (IAM) to 55 mM and incubating the solution for 40 min in darkness at room temperature (RT). A final incubation with 100 mM DTT was performed at RT before adding 2 µg of trypsin and incubating overnight at 37°C. Fractionation of samples was performed using a 3100 OFFGEL fractionator and OFFGEL kit (pH 3–10; 24-well format) (Agilent Technologies, Australia) according to the manufacturer’s protocols. Briefly, resultant peptides were diluted in peptide-focusing buffer, 150 µL of sample was loaded into each of the 24 wells and the sample was focused in a current of 50 µA until 50 kilovolt hours was reached. Each of the fractions was collected, dried under a vacuum centrifuge and resuspended in 10 µL of 0.1% TFA before being desalted using a Zip-Tip^®^ (Merck Millipore, USA). Finally, samples were dried again under a vacuum centrifuge and stored at −80°C until use.

### Mass spectrometry

Due to limited availability of material because of the difficulty in maintaining *S*. *haematobium* in mice, different numbers of replicates were run for each sample. A total of 48 offgel samples from 2 different replicates (24 samples each) were run for adult ES, 48 offgel samples from 2 different replicates (24 samples each) for adult tegument proteins, 24 offgel samples (from 1 replicate) for egg ES and 2 samples (from 2 replicates) for SEA.

All samples were analyzed by LC-MS/MS on a Shimadzu Prominance Nano HPLC coupled to a Triple Tof 5600 mass spectrometer (ABSCIEX) using a nano electrospray ion source. Samples were resuspended in 30 µl of solvent A (0.1% formic acid (aq)) and fifteen µl was injected onto a 50 mm x 300 µm C18 trap column (Agilent Technologies) at a rate of 60 µl/min. The samples were desalted on the trap column for 6 minutes using solvent A at the same rate, and the trap column was then placed in-line with an analytical 150 mm x 100 µm 300SBC18, 3.5 µm nano HPLC column (Agilent Technologies) for mass spectrometry analysis. Peptides were eluted in a linear gradient of 2-40% solvent B (90/10 acetonitrile/ 0.1% formic acid (aq)) over 80 min at 500 nL/min flow rate after 6 minutes of wash at 2% solvent B and followed by a steeper gradient from 40% to 80% solvent B in 10 min. Solvent B was held at 80% for 5 min for washing the column and returned to 2% solvent B for equilibration prior to the next sample injection. Mass spectrometer settings were: ionspray voltage = 2,200V, declustering potential (DP) = 100V, curtain gas flow = 25, nebuliser gas 1 (GS1) = 12 and interface heater = 150°C. The mass spectrometer acquired 250 ms full scan TOF-MS data followed by 20 by 250 ms full scan product ion data in an Information Dependent Acquisition (IDA) mode. Full scan TOF-MS data was acquired over the mass range 300-1600 and for product ion ms/ms 80-1600. Ions observed in the TOF-MS scan exceeding a threshold of 150 counts and a charge state of +2 to +5 were set to trigger the acquisition of product ion, ms/ms spectra of the resultant 20 most intense ions. The data was acquired using Analyst TF 1.6.1 (ABSCIEX).

### Database search and protein identification

Peak lists obtained from MS/MS spectra were identified using a combination of five search engines in SearchGUI version v3.3.3 [38] since it has been shown that combining multiple search engines increases the confidence of identified peptide spectrum matches (PSMs), distinct peptide sequences and proteins [39]. The search engines used were: X!Tandem version X! Tandem Vengeance (2015.12.15.2) [40], MS-GF+ version Release (v2018.04.09) [41], Comet version 2018.01 rev. 0 [42], MyriMatch version 2.2.140 [43] and Tide [44].

Protein identification was conducted against a concatenated target/decoy version of the *S*. *haematobium* proteome downloaded from Parasite Wormbase (version of 2017-05-WormBase - www.parasite.wormbase.org, 11,140 sequences) and concatenated to the common repository of adventitious proteins (cRAP, https://www.thegpm.org/crap/, 116 sequences) and the sequence of an antigen ortholog to *S*. *mansoni* TSP-2 that was obtained from GenBank (MK238557) (total of 11,257 (target) sequences). The decoy sequences were created by reversing the target sequences in SearchGUI. The identification settings were as follows: Trypsin specific with a maximum of 2 missed cleavages, 10.0 ppm as MS1 and 0.2 Da as MS2 tolerances; fixed modifications: Carbamidomethylation of C (+57.021464 Da), variable modifications: Deamidation of N (+0.984016 Da), Deamidation of Q (+0.984016 Da), Oxidation of M (+15.994915 Da), fixed modifications during refinement procedure: Carbamidomethylation of C (+57.021464 Da), variable modifications during refinement procedure: Acetylation of protein N-term (+42.010565 Da), Pyrolidone from E (−18.010565 Da), Pyrolidone from Q (−17.026549 Da), Pyrolidone from carbamidomethylated C (−17.026549 Da). All algorithm specific settings are listed in the Certificate of Analysis available in Supplementary Tables 1-4.

Peptides and proteins were inferred from the spectrum identification results using PeptideShaker version 1.16.27 [45]. PSMs, peptides and proteins were validated at a 1.0% False Discovery Rate (FDR) estimated using the decoy hit distribution. All validation thresholds are listed in the Certificate of Analysis available in the Supplementary Tables 1-4. Mass spectrometry data along with the identification results have been deposited to the ProteomeXchange Consortium via the PRIDE partner repository [46] with the dataset identifiers PXD011137 and 10.6019/PXD011137.

### Bioinformatic analysis of proteomic sequence data

The programs Blast2GO v5.2 [47] and HMMER v3.1b1 [48] were used to classify the proteins according to GO categories and Pfam domains respectively. Pfam domains were detected at the P<0.01 threshold for the HMMER software. ReviGO was used to visualise GO terms using semantic similarity-based scatterplots [49]. The UpSetR package (v. 1.3.3) [50] was used to visualise the intersections of proteins between samples by producing UpSetR plots in R.

### Enzyme-linked immunosorbent assay with human urine

The urine of infected individuals (n= 98) from an area in Zimbabwe mono-endemic for *S*. *haematobium* infection was analyzed by ELISA using all *S*. *haematobium* protein preparations described earlier. Urine from Australian volunteer donors that had never travelled to schistosomiasis endemic areas was used as a negative control (n= 14).

Polystyrene microtiter plates (Greiner Bio-One, Austria) were coated overnight at 4°C with 50 µl/well of a 5 µg/ml solution of *S*. *haematobium* adult ES, adult tegument, SEA or egg ES in 0.1 M carbonate coating buffer, pH 9.6. The plates were washed three times with PBS/0.05% Tween20 (PBST) and blocked for two hours at 37°C using 5% skimmed milk in PBST, followed by three wash steps in PBST for 15 min each. Plates were then incubated with 50 µl of urine (diluted 1:5 in PBS) and incubated at 37°C for 1.5 h followed by 3 washes using PBST. Fifty µl of HRP-conjugated polyclonal anti-human IgG (Sigma-Aldrich) was added at dilution of 1:5,000 and incubated for 1 hour at 37°C. Finally, plates were washed 3× with PBST and incubated with 3,3’,5,5;-tetramethylbenzidine (TMB, Thermo Fisher Scientific, USA) for 10 min at RT in the dark. The reaction was stopped by adding 3 M HCl and absorbance read at 450 nm using a POLARstar Omega spectrophotometer (BMG Labtech, Australia).

### Statistical analysis

GraphPad Prism 7.0 was used for statistical analyses. Differences in antibody titers were analysed using the non-parametric Kruskal-Wallis test with Dunn’s multiple comparisons test. Receiver Operating Characteristic (ROC) curves were used to calculate sensitivity, specificity and the area under the curve (AUC). The AUC is a global measure of diagnostic accuracy and indicates the probability of accurately identifying true positives, where a value of 0 indicates a perfectly inaccurate test and a value of 1 reflects a perfectly accurate test [51].

## Results

### *Schistosoma haematobium* tissue proteomes involved at the host-parasite interface

The proteomes from different parasite extracts involved in host-parasite interactions, such as the adult ES and tegument proteins as well as the egg ES and somatic (SEA) proteins were characterised by LC-MS/MS. To limit the number of false identifications we appended 116 sequences of common contaminants to the *S*. *haematobium* protein database (Bioproject PRJNA78265, www.parasite.wormbase.org/), and only proteins identified with ≥2 peptides were considered as positively identified. The extract with the highest number of identified proteins was the adult tegument (662, Supplementary Table 5), followed by adult ES (239, Supplementary Table 6), SEA (210, Supplementary Table 7) and egg ES (138, Supplementary Table 8). The adult tegument sample also had the highest number of unique (only present in this extract) proteins (430), followed by SEA (61), adult ES (48) and egg ES (19) (Fig 1A). Furthermore, adult tegument and adult ES proteins had 85 proteins in common, while only 57 proteins were commonly identified in all datasets (Fig 1A).

**Figure 1.**
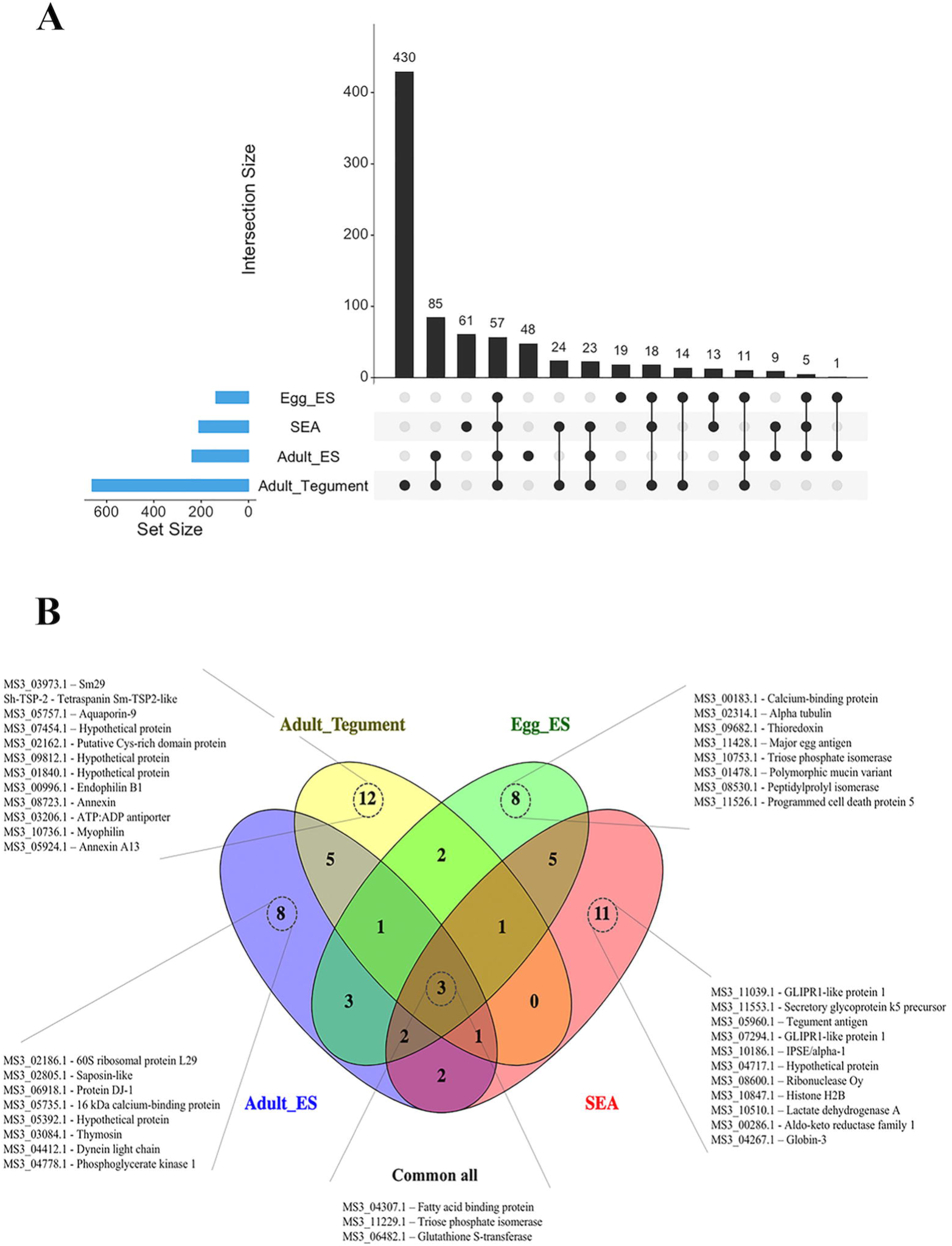
Proteins identified in the different *Schistosoma haematobium* proteomes. (A) The numbers and intersections of the identified proteins from different *Schistosoma haematobium* proteomes was visualised using an Upset plot. Connected dots display shared proteins between datasets, and the total number of proteins identified in a particular dataset is indicated in the set size. (B) Venn diagram representing the intersection of the top 25 most abundant proteins from each dataset based on spectrum counting.

The top 25 most abundant proteins in each extract selected based on the spectral count provided a similar profile: 12 proteins were uniquely identified in adult tegument extract, followed by SEA (11) and adult ES and egg ES (8 each) (Fig 1B). Only 3 proteins were commonly identified in all datasets (MS3_04307.1, fatty acid binding protein - FABP; MS3_11229.1, triose phosphate isomerase; and MS3_06482.1, glutathione S-transferase - GST) Unique proteins identified from the adult tegument included three hypothetical proteins (MS3_07454.1, MS3_09812.1 and MS3_01840.1), one tetraspanin (*Sh*-TSP-2) and two annexins (MS3_08723.1 and MS3_05924.1), among others (Fig 1B). Among the proteins uniquely identified in SEA, we found two proteins of the glioma pathogenesis related-1 (GLIPR1) subfamily (MS3_11039.1 and MS3_07294.1), IPSE/alpha-1 protein (MS3_10186.1), as well as several enzymes (MS3_08600.1, MS3_10510.1 and MS3_00286.1) (Fig 1B). A calcium binding protein (MS3_00183.1), alpha-tubulin (MS3_02314.1), thioredoxin (MS3_09682.1) and a major egg antigen (MS3_11428.1) were the most highly represented proteins uniquely identified in egg ES, while uniquely identified proteins in adult ES included saposin (MS3_02805.1) and thymosin (MS3_03084.1).

### Protein families and functions in the different proteomes of *Schistosoma haematobium*

A Pfam analysis was performed on the different proteomes from *S*. *haematobium* and the top 25 most represented protein families for each dataset were visualized in a heatmap (Fig 2). Five different protein domains (three different EF-hand-like domains, a tetratricopeptide repeat and an AAA domain) were highly represented in all proteomes, while three domains were exclusively found in proteins identified in the tegument of *S*. *haematobium* (ADP-ribosylation, Ras family and Ras of Complex Roc) (Fig 2). Similarly, one domain was found in proteins exclusively identified in the egg ES (KH domain), while other cytoskeletal and redox domains (e.g. redoxin, tropomyosin and glutathione S-transferase) were more abundant in proteins from egg ES and SEA. Interestingly, two immunoglobulin domains were highly represented in the proteins identified from the adult ES and also found in the adult tegument but not present in any of the egg proteomes (Fig 2).

**Figure 2.**
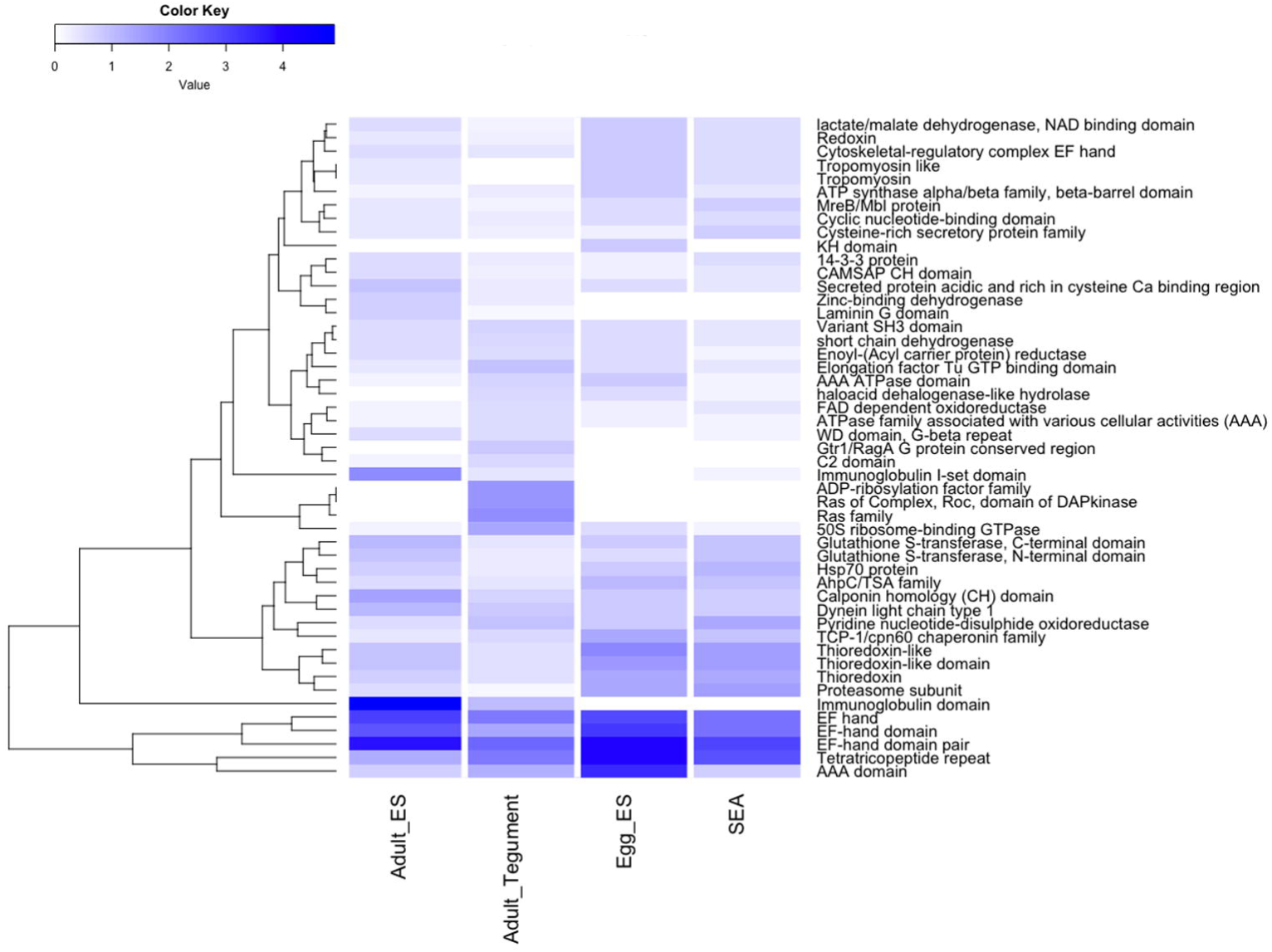
Pfam analysis. The top 25 most represented protein families from every proteome analysed were visualised using a heatmap. Values represent the abundance (in percentage) of each protein family relative to the total number of protein families present in each proteome.

The GO analysis was performed using Blast2GO [47] and biological processes and molecular functions plotted using ReviGO [49]. Several metabolic processes were highly represented in adult ES (Supplementary Table 9), together with “gluconeogenesis” and “proteolysis” (Fig 3A). An “oxidation-reduction process” was common between adult ES and tegument proteomes, while the tegument proteins were also involved in “phosphorylation” (Fig 3C, Supplementary Table 10). Regarding molecular functions inferred from adult ES and tegument proteins, both datasets were enriched in “oxidoreductase activity”, “ATP binding”, “transferase activity” and “cytoskeletal protein binding”, although presenting different scores and frequencies (Fig 3C, D).

**Figure 3.**
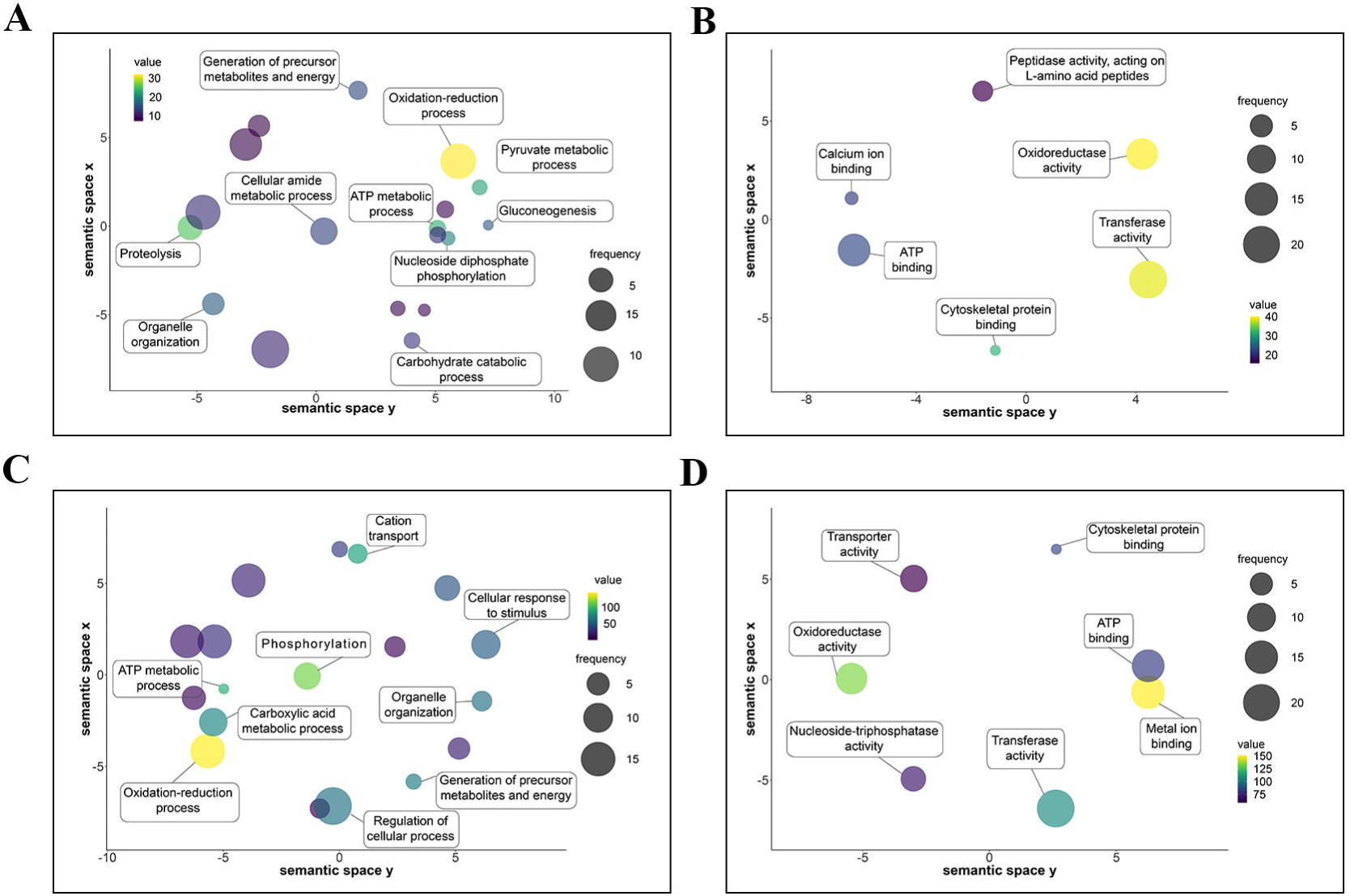
Gene ontology analysis of the adult excretory/secretory (ES) and tegument protein preparations from *Schistosoma haematobium*. Gene ontologies of proteins from *S*. *haematobium* adult ES (A, B) and tegument (C, D) were ranked by nodescore (Blast2GO) and plotted using ReviGO. Figure shows Biological Processes (A, C) and Molecular Function (B, D). Semantically similar GO terms plot close together, circle size denotes the frequency of the GO term from the underlying database, and increasing heatmap score signifies increasing nodescore from Blast2GO.

Both egg ES and SEA had distinct profiles with regards to the terms associated with biological process (Supplementary Tables 11 and 12, respectively). Despite both proteomes contain proteins involved in “gluconeogenesis” and “proteolysis”, they had different associated metabolic processes (Fig 4A, C). Two molecular function terms were similar among all proteomes (“oxidoreductase activity”, “ATP binding”), while “protein binding” was exclusively found in egg ES and SEA (Fig 4B, D). In addition, the egg ES proteome was enriched in proteins involved in “kinase activity” and “calcium ion binding”, while the molecular functions in SEA proteins related to “transferase activity” and “metal ion binding” (Fig 4B, D).

**Figure 4.**
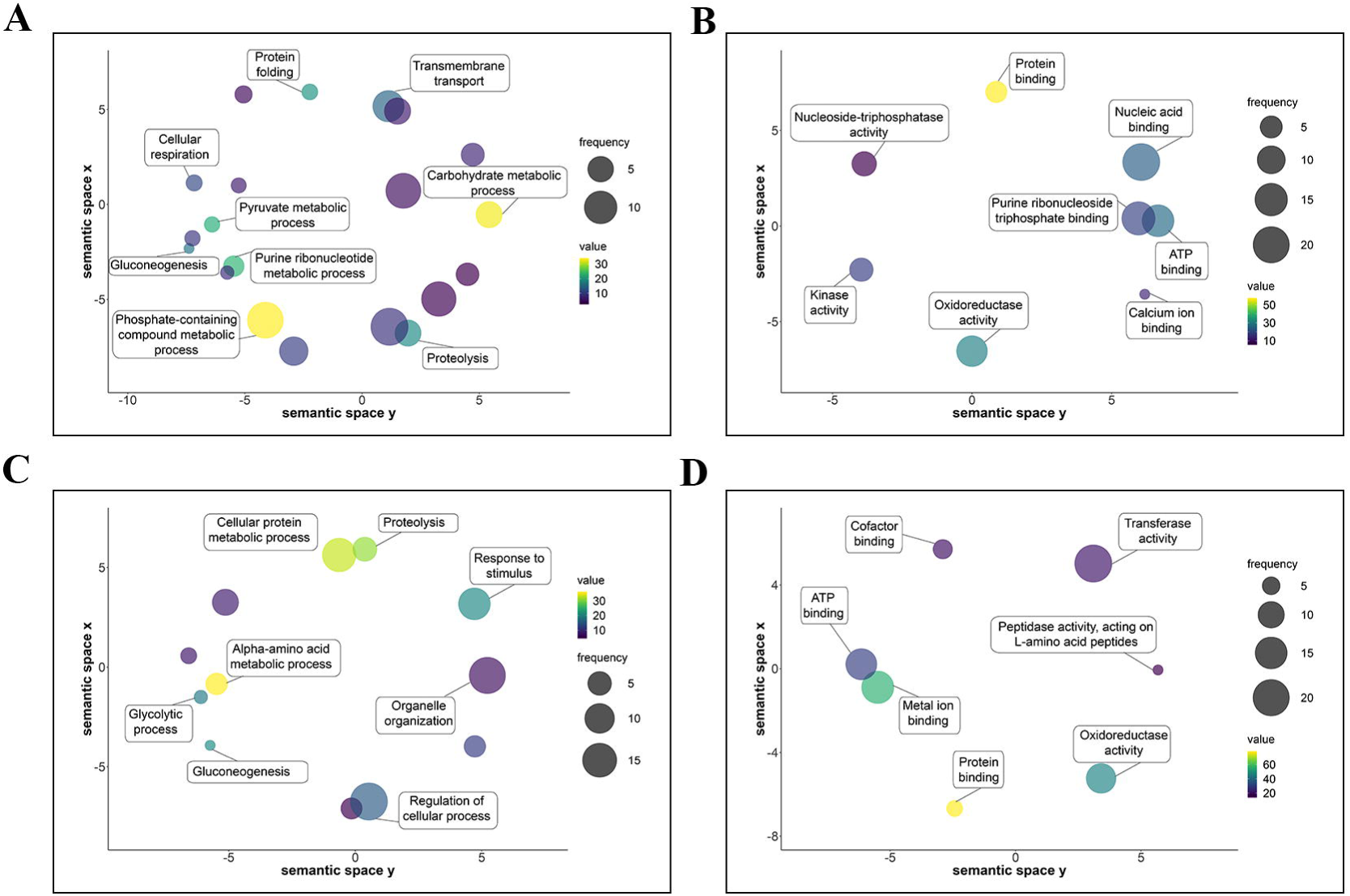
Gene ontology analysis of the egg excretory/secretory (ES) and soluble egg antigens (SEA) from *Schistosoma haematobium*. Gene ontologies of proteins from *S*. *haematobium* egg ES (A, B) and SEA (C, D) were ranked by nodescore (Blast2GO) and plotted using ReviGO. Figure shows Biological Processes (A, C) and Molecular Function (B, D). Semantically similar GO terms plot close together, circle size denotes the frequency of the GO term from the underlying database, and increasing heatmap score signifies increasing nodescore from Blast2GO.

### *Schistosoma haematobium* secreted and tegument proteins have potential as diagnostic markers of infection

Using the different *S*. *haematobium* proteomes characterized in this study, we analyzed antibody responses in the urine of *S*. *haematobium*-infected individuals (Fig 5A-D) to the various parasite extracts. Furthermore, the AUC generated from the ROC curves were used to determine the sensitivity and specificity of each antibody response, and the predictive value of infection (Fig 6A-D). Antibodies to all extracts were significantly reactive in the urine of all subjects with a high egg count (>50 eggs/ml) compared to non-endemic negative controls with the most significant reactivity being to adult fluke ES and SEA (p<0.0001), although adult tegument and egg ES samples were also highly reactive (p<0.001). Antibodies in the urine of subjects with a medium intensity infection (11-50 eggs/ml) significantly reacted to all extracts, compared to non-endemic negative controls, with the most significant reactivity being to adult ES and tegument extracts (p<0.01). Interestingly, the antibody levels in the urine of subjects with low intensity infection (0.3-10 eggs/ml) were significantly elevated only for adult ES (p<0.05) and not for other extracts tested, while subjects with a very low level of infection (egg-negative by microscopy but CAA-positive) had significantly elevated antibody levels to only adult tegument and SEA extracts (p<0.05).

**Figure 5.**
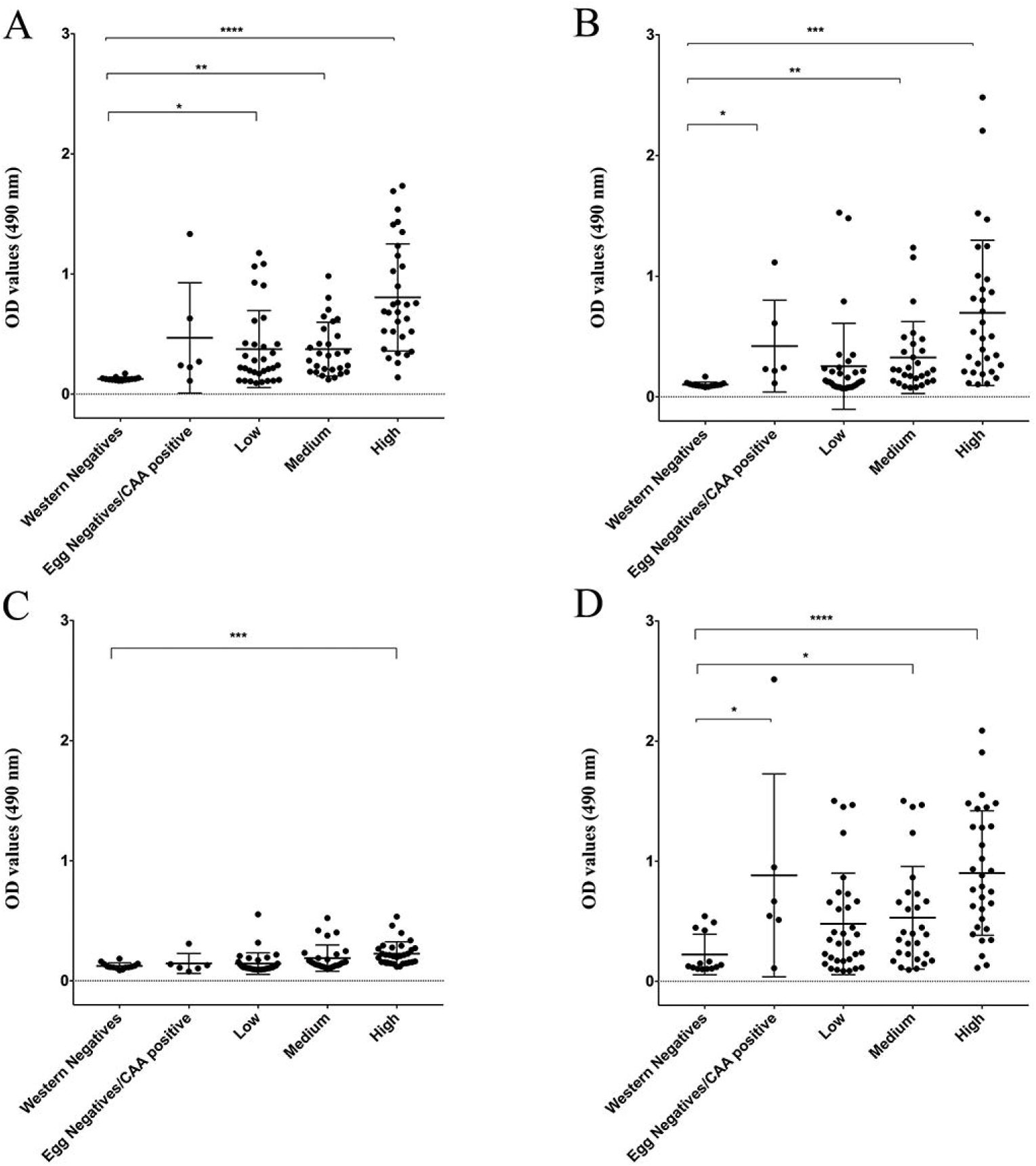
Urine antibody levels against different *Schistosoma haematobium* antigen preparations. OD values determined by ELISA using urine from individuals with different infection intensities based on egg counts (high, medium, low and egg negative/CAA positive) from an endemic area of Zimbabwe against different proteomes from *S*. *haematobium*. A control group of urine from non-endemic individuals who had never been infected with *S*. *haematobium* was included. Adult excretory/secretory (ES) products (A), tegument proteins (B), egg ES (C) and soluble egg antigen (D). Statistical analysis was performed using a non-parametric Kruskal-Wallis test with multiple comparisons by Dunn’s post-test. * *P* □< □0.05, ** *P*□ <□ 0.01, *** *P*□ <□ 0.001, **** *P*□ < □0.0001

**Figure 6.**
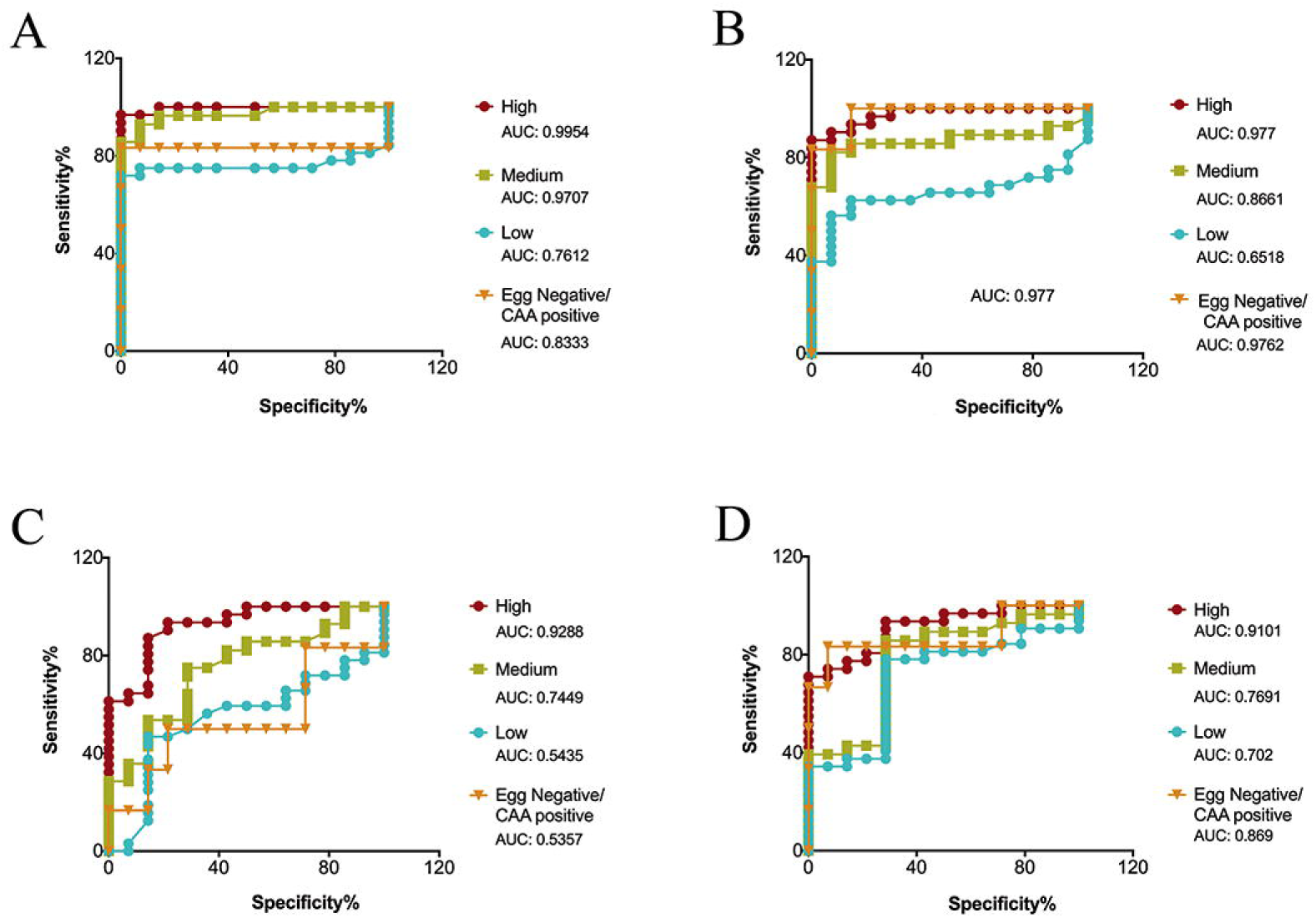
Receiver operating characteristic (ROC) curves. Areas under the ROC curve determine the predicative performance of adult excretory/secretory (ES) products (A), tegument proteins (B), egg ES products (C) and soluble egg antigen (SEA) (D) to detect antibodies in the urine of individuals with differing intensities (high, medium, low and egg negative/CAA positive) of *Schistosoma haematobium* infection. A control group of urine from non-endemic individuals who had never been infected with *S*. *haematobium* was included.

The highest predictive value of infection in subjects with a high egg count was generated by the antibody response to adult ES (AUC 0.9954), followed by adult tegument (0.977), egg ES (0.9288) and SEA (0.9101) antibody responses. In the case of subjects with a medium egg count, the highest predictive value of infection was also observed with adult ES antibody responses (0.9707), followed by adult tegument (0.8661), SEA (0.7691) and egg ES (0.7449). This was again the case for subjects with a low egg count; the predictive value of infection for adult ES, adult tegument, SEA and egg ES antibody responses being 0.7612, 0.6518, 0.702 and 0.5435, respectively. Interestingly, the highest predictive value of infection in microscopy egg negative but CAA positive subjects was generated with the antibody response to adult tegument extract (0.9762), followed by SEA (0.869), adult ES (0.8333) and egg ES (0.5357).

## Discussion

Despite accounting for almost two-thirds of all cases of schistosomiasis [52], *S*. *haematobium* remains the most neglected schistosome of medical importance in terms of laboratory research effort, partly due to the lack of proteomic and genomic information (which underpin knowledge of the parasite’s biological and pathogenic processes) [8]. Several versions of the *S*. *mansoni* genome have been published [53, 54], as well as numerous high-throughput proteomes from different life stages [22, 24–26, 55, 56]; however, only one version of the genome and very few proteins have been characterised for the causative agent of urinary schistosomiasis [57]. Indeed, a thorough examination of the *S*. *haematobium* genome failed to detect the presence of a homologue of *Sm*-TSP-2 in this parasite, despite this protein being found in the proteomic datasets described herein, its abundance in *S*. *mansoni* proteomes [58] and the presence of numerous homologs in *S*. *japonicum* [59]. *Sm*-TSP-2 has proven to be one of the most effective vaccine candidates against schistosomiasis [60, 61] and has successfully undergone a phase I clinical trial [62]; confirmation of the existence of a *S*. *haematobium* orthologue is important, therefore, to determine whether a similar (or existing) vaccination strategy might be effective against this species of schistosome. This example highlights the importance of the availability of proteomic data to complement and, as is being increasingly reported, re-annotate existing genomes to provide a comprehensive picture of the makeup of an organism [86].

Proteins at the interface between the host and the parasite are believed to play a key role in host immune system modulation and parasite survival [63] and so characterisation of these molecules is desirable to further our knowledge of parasite biology and pathogenesis and intervention targets. For example, the proteins secreted and located on/in the tegument of the blood-dwelling adult fluke are immune-accessible, and egg ES molecules are involved in the generation of a tumorigenic environment [13] and can be studied to gain insight into host pathogenesis.

A total of 57 proteins were commonly identified in all proteomic datasets generated. Among these, MS3_04307.1 (FABP), MS3_11229.1 (triose phosphate isomerase) and MS3 06482 (GST) were highly abundant. All proteomes were enriched in three main protein families: AAA domains, EF-hand like domains and tetratricopeptide repeat (TPR) domains. The high abundance of proteins containing these domains suggests a vital role for these motifs in parasite development and/or survival. While AAA domain-containing proteins such as dyneins participate in a variety of functions and are ubiquitously present in many organisms [64], EF-hand domain proteins are expressed almost uniquely by parasitic worms (highly abundant in trematodes), and have been suggested to be an effective drug target [65]. TPR is a structural repeat found in diverse proteins. In *Schistosoma mekongi* it is present within an O-glycosyltransferase that is abundantly expressed by female flukes [66], while in *S*. *japonicum*, the protein SJCHGC06661 was the highest immunoreactive protein in a protein array screened against the serum from *S*. *japonicum* infected patients. This protein contains six TPR domains, interacts with the heat shock protein (HSP) complex and has been suggested as a potential drug or vaccine target [67].

The most complex mixture of proteins was identified in the adult tegument sample (662 proteins). Previous studies on *S*. *mansoni* schistosomula and *S*. *japonicum* adult flukes identified 450 and 360 proteins, respectively [22, 68], highlighting the complex composition of this structure, probably due to an adaptive response to living in the harsh environment of the host circulatory system. The molecular functions associated with these proteins correspond to the functions observed in other studies [68]. For instance, proteins involved in several binding functions (the highest represented function in the proteins identified in the *S*. *japonicum* adult tegument [68]) such as metal ion binding, cytoskeletal protein binding and ATP binding were highly represented in this study. Several molecular functions associated with protection against oxygen-reactive species and redox systems such as oxidoreductase activity and transferase activity were also highly represented, as expected in the tegument of a parasitic helminth which is under constant immune threat [69]. Among the most abundant proteins present in the tegument of *S*. *haematobium* adult flukes, we identified a putative GPI-anchored protein similar to Sm29 (MS3_03973.1), a tetraspanin similar to *Sm*-TSP-2 (*Sh*-TSP-2), an aquaporin (MS3_05757.1), as well as two different annexins (MS3_08723.1 and MS3_05924.1) and several hypothetical proteins (MS3_07454.1, MS3_09812.1 and MS3_01840.1). Sm29 has been shown to be an immunoregulatory molecule able to control inflammatory mucosal diseases with both Th1 and Th2 immune response profiles [70] and can interact with CD59, an inhibitor of the membrane attack complex (MAC), which could contribute to immune evasion [71]. Tetraspanins have a role in maintaining the structure of the tegument and providing stability [72]. Interestingly, Sm29 and *Sm*-TSP-2, as well as the above-mentioned GST and FABP, are important vaccine candidates [27]. Characterising the rest of the identified proteins, particularly the hypothetical proteins, will improve the rational selection of vaccine candidates for *S*. *haematobium* infection.

Using an older version of the *S*. *haematobium* proteome present in WormbaseParasite, only 379 proteins were predicted to be secreted by *S*. *haematobium*, making it the parasitic helminth with the lowest number of predicted secreted proteins of the 44 species assessed [73]. In our analysis, we found 239 proteins in the adult ES products, which accounts for more than 63% of the total predicted secretome. The most abundant domains identified by Cuesta-Astroz in the predicted secretome were PF00053 (Laminin-EGF), which was not highly represented in our analysis, and PF13895 (Immunoglobulin domain), which was one of the most abundant families in the adult ES dataset. We also identified the Immunoglobulin I-set domain as an abundant protein in the adult ES products, which is also one of the most commonly occurring proteins in the predicted helminth secretomes [73], and is present in cell adhesion molecules.

Egg secretions from trematodes likely result from an active secretion process of the miracidium enclosed within the egg and, in the case of *S*. *haematobium*, the proteins secreted by the egg might not only reflect the biology of the miracidium stage, but might also be of importance in the development of fibrosis and cancer. Among the top 25 most abundant proteins in all datasets, 8 proteins were uniquely identified in the egg secretions, including calcium-binding protein (MS3_00183.1), alpha-tubulin (MS3_02314.1), thioredoxin (MS3_09682.1), polymorphic mucin variant (MS3_01478.1) and the major egg antigen (MS3_11428.1). The calcium-binding protein is orthologous (90% identity) to *S*. *mansoni* calmodulin, which has been reported as an essential protein for egg development [74]. Indeed, calmodulin inhibitors can disrupt egg hatching and interrupt miracidium transformation into sporocyst [75, 76]. The polymorphic mucin variant has been identified in the miracidium of *S*. *mansoni* as having a key role in invertebrate host/parasite interactions [77], and a family of *S*. *mansoni* polymorphic mucins has been hypothesized to act as a smoke-screen blocking pattern recognition receptors, thus avoiding recognition by the host immune system [77].

Different proteins with a role in defence against oxidative stress, such as GST and thioredoxin, were abundant in the egg ES proteome; as well as two triose phosphate isomerases, which are proteins implicated in glycolysis and energy production. Interestingly, two homologs of programmed cell death protein 5 (PCDC5; MS3_11526.1) and the major egg antigen also known as Smp40 (MS3_11428.1) were also uniquely identified in the egg ES and are among the top 25 most abundant proteins. PCDC5 has a wide variety of biological functions, including programmed cell death and immune regulation [78], and a decrease in the levels of this protein has been associated with multiple types of cancers [78]. Smp40 is associated with reduced collagen deposition, decreased fibrosis, and granuloma formation inhibition [79]. Active secretion of these two molecules might reflect an effort from the parasite to minimize the potential fibrosis and cell proliferation induced by other proteins present in the egg shell.

Two GLIPR1 proteins (MS3_11039.1 and MS3_07294.1) were uniquely identified in the SEA dataset and were among the most abundant proteins based on spectrum counting. These proteins belong to the sperm-coating proteins/Tpx1/Ag5/PR-1/Sc7 (SCP/TAPS) family, and have been identified in other trematodes including *Fasciola hepatica* [80] as well as other helminths [81]. Although the exact function for these proteins is still unknown, this family of proteins is expanded in the genomes and secreted proteomes of clade IV and V nematodes [82–86] and is believed to play specific biological functions in the host including defence against host attack and determination of lifespan [87]. The presence of these proteins in the eggs of *S*. *haematobium* is intriguing and their roles in this life stage warrants exploration.

The IPSE/alpha-1 glycoprotein (MS3_ 10186.1) is a well characterized molecule from the eggs of *S*. *mansoni*. It is located in the subshell area of mature eggs [88] and is a potent driver of IL-4-mediated Th2 responses [88–91]. Furthermore, it induces a potent anti-inflammatory response by inducing regulatory B cells to produce IL-10 [89]. The *S*. *haematobium* homolog of IPSE/alpha-1 (H-IPSE) infiltrates host cells and translocate to the nucleus [19] and, interestingly, has been shown to alleviate the symptoms of chemotherapy-induced hemorrhagic cystitis [20].

With a view to providing the first steps towards the characterisation of novel molecules that could be used for immunodiagnosis of *S*. *haematobium* infection, we assessed the levels of IgG antibodies present in the urine of patients from an endemic area in Zimbabwe against the different extracts analysed in the study. Even though the presence of antibodies in serum or urine cannot be used to differentiate previous and current infections, the detection of immunoglobulins against parasites is still a useful diagnostic tool for surveillance, and can be helpful for evaluating the effectiveness of control programmes [92]. Antibody detection diagnostics can also be used to complement less sensitive point-of-care (POC) diagnostic tests, such as microscopic detection of eggs in urine, in areas of low transmission. Some efforts have been made to develop highly sensitive and specific tests for *S*. *haematobium* infection. For instance, diverse parasite proteomes such as SEA, cercarial antigen preparation (CAP) and soluble worm antigen preparation (SWAP) have been tested [93]. In this study, SEA and CAP were more reactive than SWAP, which is in accordance with previous findings [92, 94–96]. The fact that SWAP is not a good extract from a diagnostic point of view could be because it is highly abundant in intracellular proteins that will never be exposed to host antibodies. The detection of antibodies against SEA and SWAP in the urine of human subjects with schistosomiasis has been previously reported [92, 95, 96], but the presence and diagnostic utility urine antibodies to adult fluke tegument and ES have not been described. Herein, we highlight the diagnostic capability of arguably more immunologically relevant proteomes, in addition to SEA, to detect antibodies in the urine of infected individuals, and show that the adult fluke ES and tegument preparations were more reactive than SEA for all of the cohorts studied. Antibodies can be detected in urine of subjects infected with both *S*. *mansoni* and *S*. *japonicum* (where eggs are passed in the stool instead of urine), so the presence of antibodies in urine of subjects infected with *S*. *haematobium* is likely due to clearance of IgG from circulation through the kidneys as opposed to vascular leakage due to bladder pathology caused by parasite eggs.

We have provided the first comprehensive high-throughput analysis of the proteins present in different *S*. *haematobium* proteomes of importance in host/parasite interactions, thereby facilitating a snapshot of the molecular biology of the parasite and how it interacts with the host. In addition, we have identified adult fluke ES and tegument extracts as best predictors of infection when probed with antibodies from the urine of infected human subjects. This combined study of proteomic characterisation and serodiagnostic analyses provide the first steps towards the characterisation of novel molecules that could serve as tools for the control and evaluation of *S*. *haematobium* infection in Sub-Saharan Africa.

## Supporting information

## Acknowledgements

This work was supported by a program grant from the National Health and Medical Research Council (NHMRC) [program grant number 1037304] and a Senior Principal Research fellowship from NHMRC to AL (1117504). The funders had no role in the study design, data collection and analysis, decision to publish, or preparation of the manuscript. The authors thank the study participants, as well as the parents/legal guardians and their teachers in Zimbabwe for their support of this study. They are very grateful for the cooperation of the Ministry of Health and Child Welfare in Zimbabwe. For their technical support, they would like to thank the members of the Department of Biochemistry at the University of Zimbabwe.

## Author contributions

### Conceptualization

Javier Sotillo, Mark S. Pearson and Alex Loukas.

### Resources

Javier Sotillo, Gebeyaw G. Mekonnen, Luke Becker, Takafira Mduluza and Francisca Mutapi.

### Investigation

Javier Sotillo, Mark S. Pearson, Abena S Amoah, Govert van Dam and Paul L.A.M. Cortsjens.

### Formal analysis

Javier Sotillo.

### Writing - original draft

Javier Sotillo.

### Writing – review & editing

Javier Sotillo, Mark S. Pearson, Francisca Mutapi and Alex Loukas

## Supporting information Captions

**Table S1.** Certificate of analysis provided by PeptideShaker for the analysis of the *Schistosoma haematobium* adult secreted proteins.

**Table S2.** Certificate of analysis provided by PeptideShaker for the analysis of the *Schistosoma haematobium* adult tegumental proteins.

**Table S3.** Certificate of analysis provided by PeptideShaker for the analysis of the *Schistosoma haematobium* egg secreted proteins.

**Table S4.** Certificate of analysis provided by PeptideShaker for the analysis of the *Schistosoma haematobium* soluble egg antigen proteins.

**Table S5.** *Schistosoma haematobium* adult secreted proteins identified using SearchGUI and PeptideShaker.

**Table S6.** *Schistosoma haematobium* adult tegumental proteins identified using SearchGUI and PeptideShaker.

**Table S7.** *Schistosoma haematobium* egg secreted proteins identified using SearchGUI and PeptideShaker.

**Table S8.** *Schistosoma haematobium* soluble egg antigen proteins identified using SearchGUI and PeptideShaker.

**Table S9.** Annotation of the *Schistosoma haematobium* adult secreted proteins provided by Blast2GO.

**Table S10.** Annotation of the *Schistosoma haematobium* adult tegumental proteins provided by Blast2GO.

**Table S11.** Annotation of the *Schistosoma haematobium* egg secreted proteins provided by Blast2GO.

**Table S12.** Annotation of the *Schistosoma haematobium* soluble egg antigen proteins provided by Blast2GO.

